# Chromosome-scale genome assembly of the tropical abalone (*Haliotis asinina*)

**DOI:** 10.1101/2024.04.02.587722

**Authors:** Roy Barkan, Ira Cooke, Sue-Ann Watson, Sally C. Y. Lau, Jan M. Strugnell

## Abstract

Abalone (family Haliotidae) are an ecologically and economically significant group of marine gastropods that can be found in tropical and temperate waters. To date, only a few *Haliotis* genomes are available, all belonging to temperate species. Here, we provide the first reference genome of the tropical abalone *Haliotis asinina*. The combination of PacBio long-read HiFi sequencing and Dovetail’s Omni-C sequencing allowed the chromosome-level assembly of this genome, while PacBio Isoform sequencing across five tissue types enabled the construction of high-quality gene models. This assembly resulted in 16 pseudo-chromosomes spanning over 1.12 Gb (98.1% of total scaffolds length), N50 of 67.09 Mb, the longest scaffold length of 105.96 Mb, and a BUSCO completeness score of 97.6%. This study identified 25,422 protein-coding genes and 61,149 transcripts. In an era of climate change and ocean warming, this genome of a heat-tolerant species can be used for comparative genomics with a focus on thermal resistance. This high-quality reference genome of *H. asinina* is a valuable resource for aquaculture, fisheries, and ecological studies.

## Background & Summary

Abalone (*Haliotis*) are a genus of marine herbivorous gastropods found in tropical and temperate coastal waters on every continent except for the Pacific coast of South America and the Atlantic coast of North America^1^. In addition to their ecological, historical, and cultural importance^2–4^, abalone are a highly prized seafood product that underpins valuable wild-harvest and aquaculture industries in many countries^5,6^. There has been a significant decrease in wild populations of abalone largely due to illegal harvesting, pollution, climate change and disease^5,7,8^. As a result, many species of abalone are recognized to be at risk – the IUCN Red List™ lists 44% of abalone species as being threatened with extinction^9,10^.

Whether in the wild or in aquaculture, abalone are also at risk due to ocean warming and extreme environmental events^11,12^. In the summer of 2011, between early February and early March, wild Roe’s abalone stocks suffered significant, if not total, mortality around Kalbarri, Western Australia^13^. Similarly, the 2016 mortality event of wild abalone near the coast of Tasmania, Australia, led to smaller catches and reduced quotas^14^. These events have great economic impacts on abalone fisheries, resulting in a significant decrease in production and loss of income. The increase in abalone aquaculture and the concerns for wild populations worldwide have motivated researchers to apply omics tools to provide genetic resources, improve knowledge regarding this genus, and ultimately aid production and conservation. To date, the great majority of the genetic resources available for abalone are of temperate species^15–20^. No reference genome for any tropical abalone species has been published to date.

The Donkey’s ear abalone, *Haliotis asinina* (Linnaeus, 1758), is the largest of the tropical abalone species. It is also the fastest-growing abalone of all abalone species^21^. This species is distributed throughout the Indo-Pacific and is highly desired as seafood, mainly in South-East Asia^22,23^. Due to its popularity, wild stocks are at risk as a result of overfishing^24^. Efforts are underway to revive *H. asinina* populations through stock enhancement and the use of marine reserves^25^. The lack of genetic data available for this species limits studies on genetic variation (between and within abalone species), development of genetic breeding programs, connectivity and genetic technologies that will assist fisheries, aquaculture and conservation strategies.

Here, we provide the first reference genome of the tropical abalone *H. asinina*. Using Pacific Biosciences of California, Inc. (PacBio) 5-base HiFi sequencing, Dovetail Genomics Omni-C approach and PacBio Isoform sequencing (Iso-Seq), we assembled and annotated the 1.14 Gb length reference genome. The total genome length was assembled into 170 scaffolds, with an N50 of 67.09Mb, L90 of 15, a BUSCO completeness score of 97.6% and a k-mer completeness of 99.5%. Over 98% of the scaffold’s length was anchored to 16 pseudo-chromosomes. The chromosome number matches the findings of the previous karyotype studies^26^. Furthermore, 40.0% of the genome was identified as repetitive sequences. A total of 25,422 protein-coding genes were predicted, including 61,149 transcripts. In addition, we used the same data to measure DNA methylation across the genome and to assemble the mitochondrial genome of *H. asinina*.

This significant resource, along with the use of omics tools (i.e., comparative genomics, transcriptomic, epigenomics and proteomics), will provide new insights regarding the evolution of abalone and genetic factors that might assist in overcoming the current and future challenges mentioned above.

## Methods

The general workflow is illustrated in **Figure 1**.

**Figure 1.**
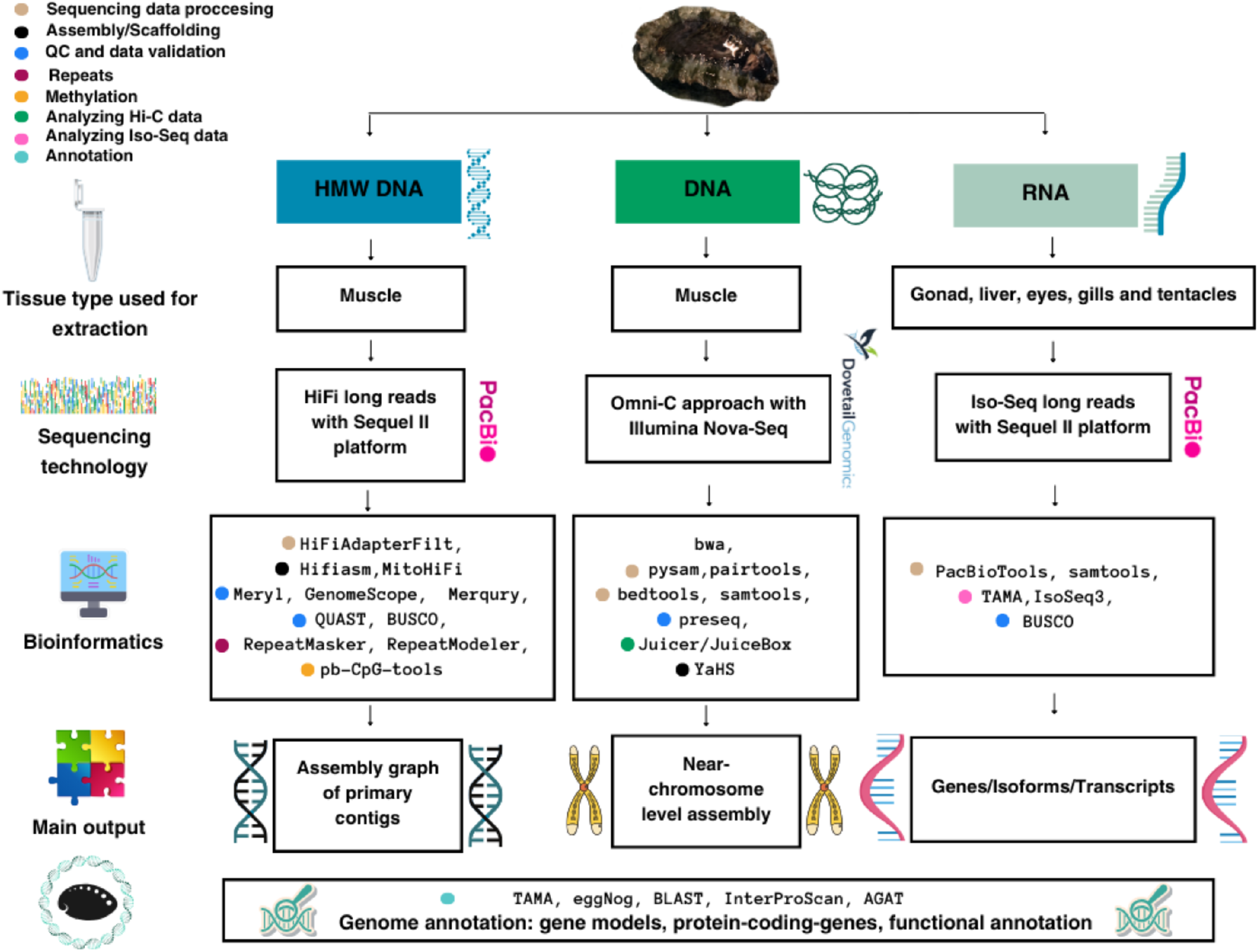
Schematic overview of the study workflow.

### Biological materials

In April 2022, *H. asinina* individuals were obtained from Arlington Reef (−16° 42’ 26.1036”S, 146° 3’ 30.4128”E) on the Great Barrier Reef, Australia, by divers from Cairns Marine Pty Ltd. Abalone were introduced into a round 100L white plastic aquaria at the Marine and Aquaculture Research Facility (MARF) at James Cook University (Townsville, Australia). High water quality was maintained during the entire period. The temperature was set to the ambient temperatures at the collection site and was recorded continuously using the facility’s automated monitoring system. Water quality parameters, including ammonia, nitrate, and nitrite, were measured using “AquaSonic” kits. Water in the aquaria was replaced every two to three days. The abalone were fed every two days using Halo abalone feed (3mm pellets) manufactured by Skretting.

### Sampling, nucleic acid extraction, library preparation and sequencing

Sampling, nucleic acid extraction, library preparation and sequencing were all performed on the same individual (described below).

Following a fasting period of 24-hours, one abalone individual (female, body length = 10.9cm, shell length = 7.4cm) was randomly selected and dissected immediately for High Molecular Weight Genomic DNA (HMW gDNA) extraction. HMW gDNA was extracted from the ∼30 mg of fresh muscle tissue using the Circulomics® Nanobind Tissue Big DNA Kit following protocol modification for *Aplysia*^27^. Library preparation and sequencing were performed by the Australian Genome Research Facility (AGRF) according to PacBio protocols. Sequencing was performed using a single SMRT Cell and the PacBio Sequel II (specifically, 5-base HiFi sequencing) with seq polymerase version 2.2 and seq primer v5. Movie time was 30hrs and 120pM SMRTcell loading. This resulted in 36.2 GB of data with 2.62M (million) high-quality reads (**Table *1***).

**Table 1.** Basic statistics of the sequencing data.

| Library type | Sequencing platform | Tissue | Reads (M) | Yield (Gb) | Read length (mean, bp) |
| --- | --- | --- | --- | --- | --- |
| PacBio (HMW gDNA) | PacBio Sequel II | Muscle | 2.62 | 36.24 | 13,825 |
| PacBio Iso-Seq (cDNA) | PacBio Sequel II | Gonad, liver, eyes, gills, tentacle | 3.13 | 5.90 | 1,886 |
| Omni-C | Illumina Novaseq X plus | Muscle | 478.26 | 144.43 | 150 |

The DNase Hi-C (Omni-C) library was prepared using the Dovetail Omni-C® Kit at AGRF according to the manufacturer’s protocol with modifications as follows: 60 mg of abalone muscle tissue was thoroughly cryo-ground using liquid nitrogen, and the chromatin was fixed with disuccinimidyl glutarate (DSG) and formaldehyde in the nucleus. After removing the cross-linking reagents, the disrupted tissue sample underwent sequential filtration through 200 μm and 50 μm cell strainers to eliminate large debris. The cross-linked chromatin was then digested *in situ* with the optimal amount of DNase I to achieve efficient chromatin digestion and, hence, generate long-range *cis* reads. Following digestion, the cells were lysed with sodium dodecyl-sulfate (SDS) to extract the chromatin fragments. Stage 3 of the library preparation - proximity ligation, was optimised (1) by reducing the recommended input lysate, thereby minimising any impurities, and (2) by increasing the intra-aggregate bridge ligation to an overnight reaction to enhance the ligation events. Briefly, optimally digested chromatin fragments were bound to Chromatin Capture Beads. Next, the chromatin ends were repaired and ligated to a biotinylated bridge adapter, followed by proximity ligation of adapter-containing ends. After proximity ligation, the crosslinks were reversed, the associated proteins were degraded, and the DNA was purified and then converted into a sequencing library using Illumina-compatible adaptors. Biotin-containing fragments were isolated using streptavidin beads prior to PCR amplification. The library was sequenced on an Illumina Novaseq X plus a platform to generate two million 2 X 150 base-pairs (bp) read pairs to assess the quality of mapping, valid cis-trans reads and complexity of the library. For chromosome-level assembly, the Omni -C library was sequenced to achieve approximately 100M 2 X 150 bp read pairs per GB of the genome size. This resulted in 144.43 GB of data, including 478.26M reads (**Table *1***).

Total RNA was extracted from five tissue types: gonad, liver, epipodial tentacle, eyes and gills. Each tissue was crushed with a sterilized, chilled pestle and mortar using 1ml of TRIzol™. Once the tissue disruption was completed, the lysate was kept at −20°C overnight. Total RNA extraction was completed using TRIzol™ Plus RNA Purification Kit (Invitrogen™) following the manufacturer’s protocol. The extracted RNA was stored at −80°C. Library prep and sequencing were performed by AGRF following PacBio protocols. Sequencing was done using the PacBio Sequel II and yielded 5.90 GB of data with 3.13M reads (**Table *1***).

### Genome assembly and scaffolding

For the assembly, we used the Hifiasm version 0.19.7^28^ haplotype-resolved de novo assembler with the PacBio HiFi adapter-free FASTA file and Hi-C partition using Omni-C data. Next, Omni-C data was used as input for Dovetail’s Omni-C workflow (https://github.com/dovetail-genomics/Omni-C). The workflow includes various tools^29–34^ for QC of the Omni-C library and generating contact maps. The primary assembly was indexed using SAMtools^32^ and the index was file was used to generate genome. Omni-C reads were aligned to a reference genome using BWA version 0.7.17^30^, and high-quality mapped reads were retained. The mapped data was used as input for pairtools version 1.0.3^31^ to identify proximity ligation events, categorize pairs by read type, insert distance, and flag and remove PCR duplicates. Juicer tools version 1.6^34^ was used to generate the HiC contact matrix and contact map. The final scaffolding step was done using YaHs version 1.1^35^, which resulted in 170 scaffolds that span over 1.14 Gb with the longest scaffold size of 105.96 Mb, N50 of 67.09 Mb and L90 of 15 (**Table 2**). The Hi-C map (**Figure 2**) suggested 16 chromosome-scale scaffolds, comprising 98.1% of the total genome size (**Table 3**).

**Table 2.** Basic statistics of *H. asinina* final genome assembly.

| Assembly parameter | *H. asinina* |
| --- | --- |
| Scaffolds | 170 |
| Total length | 1,144,142,131 |
| Largest scaffold | 105,963,769 |
| GC (%) | 40.13 |
| N50 | 67,096,918 |
| N90 | 52,850,021 |
| L50 | 7 |
| L90 | 15 |

**Table 3.** Basic statistics of the 16 pseudo-chromosomes.

| Name | Length (bp) | Percentage | Number of genes | Percentage |
| --- | --- | --- | --- | --- |
| Chr_1 | 105,963,769 | 9.26% | 2085 | 9.20% |
| Chr_2 | 101,978,182 | 8.91% | 2256 | 9.95% |
| Chr_3 | 89,013,229 | 7.78% | 1891 | 8.34% |
| Chr_4 | 83,060,790 | 7.26% | 1195 | 5.27% |
| Chr_5 | 79,603,559 | 6.96% | 1659 | 7.32% |
| Chr_6 | 71,828,453 | 6.28% | 1564 | 6.90% |
| Chr_7 | 67,096,918 | 5.86% | 1594 | 7.03% |
| Chr_8 | 66,935,639 | 5.85% | 1656 | 7.31% |
| Chr_9 | 66,300,187 | 5.79% | 1421 | 6.27% |
| Chr_10 | 59,771,656 | 5.22% | 1306 | 5.76% |
| Chr_11 | 58,626,679 | 5.12% | 907 | 4.00% |
| Chr_12 | 58,286,999 | 5.09% | 1264 | 5.58% |
| Chr_13 | 55,744,664 | 4.87% | 761 | 3.36% |
| Chr_14 | 55,702,281 | 4.87% | 1104 | 4.87% |
| Chr_15 | 52,850,021 | 4.62% | 1088 | 4.80% |
| Chr_16 | 49,714,875 | 4.35% | 912 | 4.02% |
| **Total** | **1,122,477,901** | **98.11%** | **22663** | **89.08%** |
| **Unplaced (153 scaffolds)** | **21,606,066** | **1.89%** | **2779** | **11.91%** |
| **Chr_159 (Mitochondrial)** | **17,450** | **-** | **37** | **-** |

**Figure 2.**
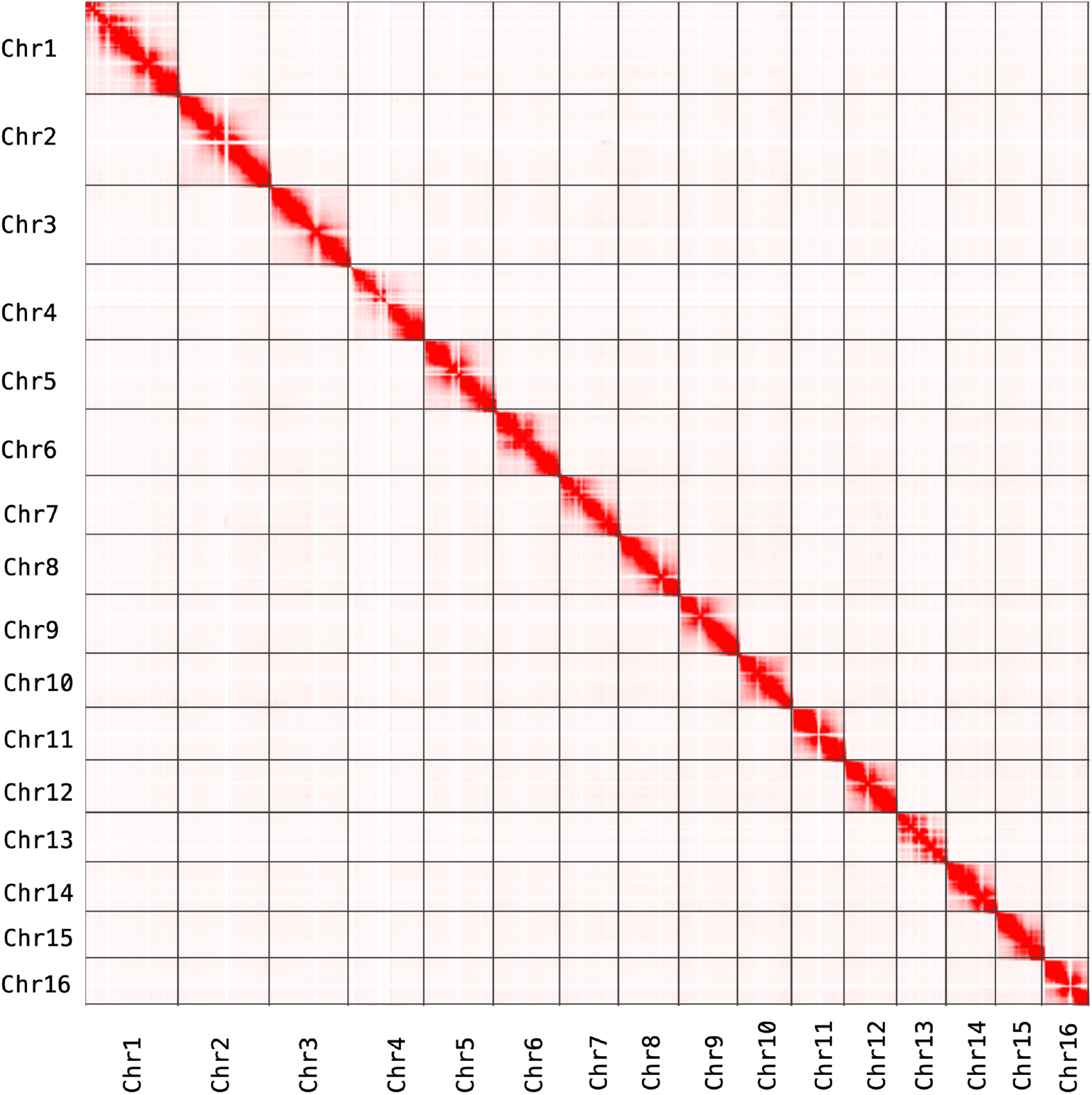
Hi-C heatmap. Pairwise interactions between pairs of chromosomes throughout H. asinina genome assembly.

### Methylation calling

High-quality reads produced using PacBio 5-base HiFi sequencing were used for CpG methylation calling across the genome assembly. Primrose version 1.3.0 (https://github.com/mattoslmp/primrose), a tool that predicts 5-methylcytosine (5mC) in HiFI reads, was used to add MM and ML tags (SAM tags that represent base modifications/methylation and base modification probabilities, respectively). The reads, which included the MM and ML tags, were aligned to the assembly using pbmm2 version 1.13.1 (https://github.com/PacificBiosciences/pbmm2), a minimap2^36,37^ SMRT wrapper for PacBio data. Then, the aligned_bam_to_cpg_scores tool provided in pb-CpG-tools version 2.3.2 (https://github.com/PacificBiosciences/pb-CpG-tools) was used to generate CpG site methylation probabilities. Then, high probability (>95%) methylation site density was calculated across the entire genome (**Figure 3**).

**Figure 3.**
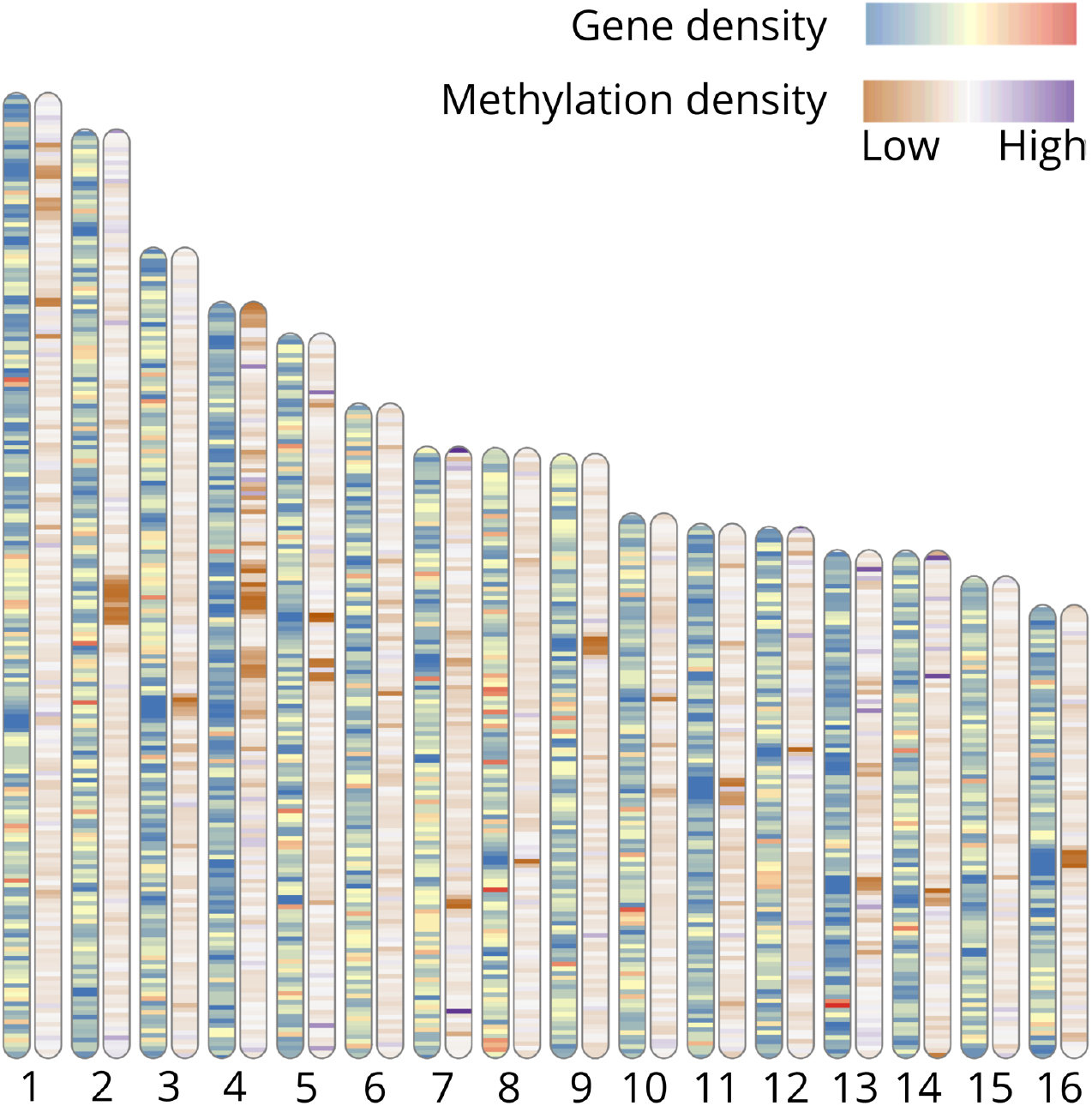
Chromosome ideogram. Each pair of ideograms represents one of the sixteen chromosomes in the *H. asinina* genome. The numbers at the bottom of each ideogram represent the number of the chromosome (i.e. 1=chr1). The inner heatmap in the left ideogram of each pair represents the gene density (bin size=0.5Mb). The inner heatmap in the right ideogram of each pair represents the methylation density (bin size=0.5Mb, probability cut-off of 95%). The top colour scale represents the gene density, and the lower scale represents the methylation density.

### Mitochondrial genome assembly

MitoHiFi version 3.0.1^38^, a pipeline for mitochondrial genome assembly from PacBio HiFi reads (or the assembled contigs/scaffolds), was used with the default annotation tools – MitoFinder version 1.4.1^39^ and ARWEN^40^. Scaffold_159 corresponds to the mitochondrial genome (17450 bp in length), including all 37 identified mitochondrial genes, with no frameshifts and high probability (>96%).

### Repetitive sequence identification

RepeatModeler version 2.0.5^41^ and RepeatMasker version 4.1.5^42^ were used to screen the *H. asinina* genome assembly for *de novo* identification of transposable elements (TEs) and classification of repeated and low complexity sequences (**Table 4**). The proportion of repeated elements in *H. asinina* genome was 38.42%, half of which were classified as unknown (19.71%). Retroelements (Class I) comprised 13.25%, DNA transposons (Class II) were 5.37%, and 1.30% were simple repeats. The proportion of repeats found in *H. asinina* genome is relatively similar to other abalone species^17,43–45^ and other marine invertebrates^46,47^ such as *Aplysia californica*^48^ and *Crassostrea virginica*^49^.

**Table 4.**
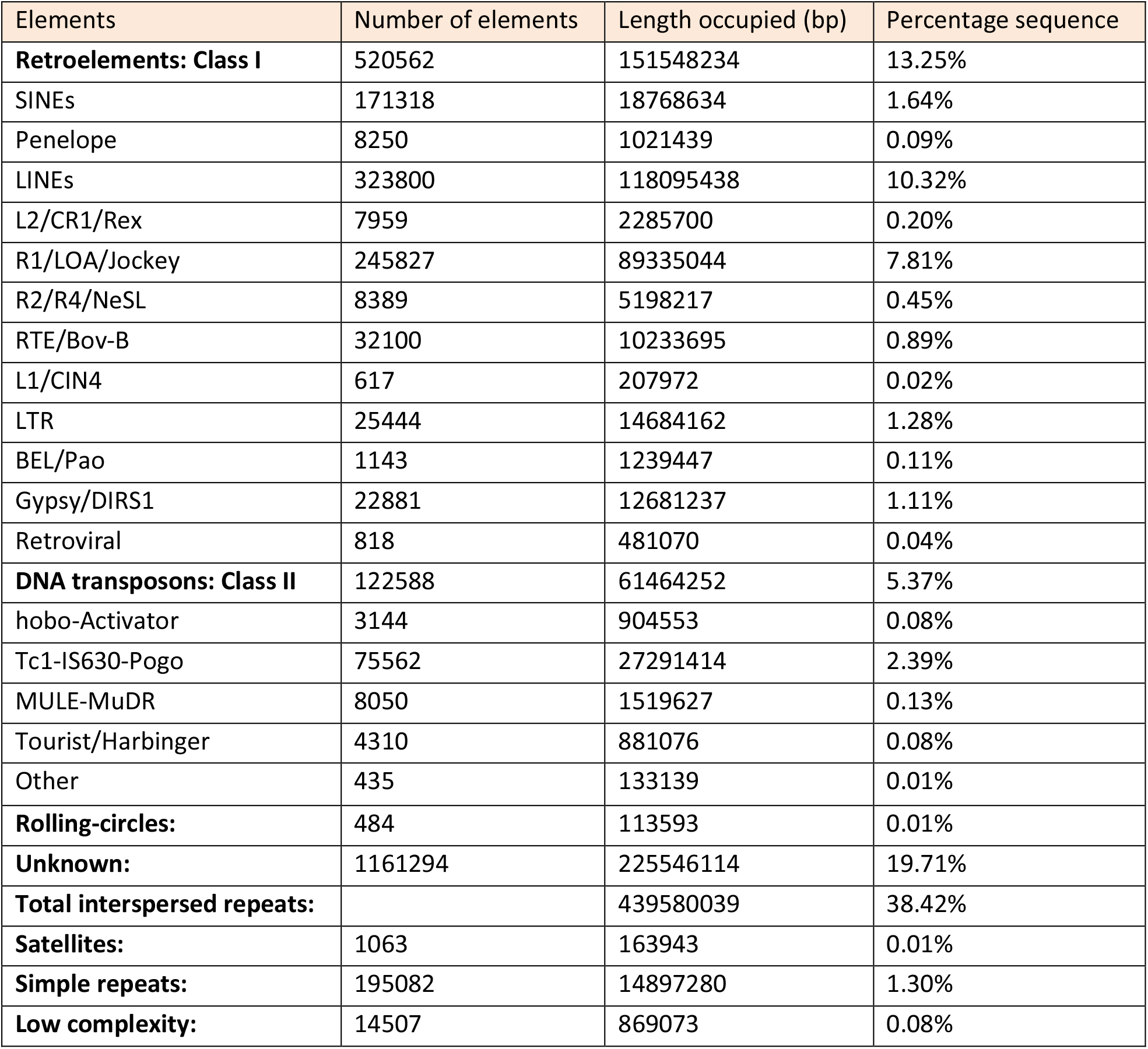
Summary of repetitive elements in the genome assembly of *H. asinina*.

### Gene prediction and functional annotation

Gene prediction was performed on a version of the genome that was soft-masked for repeats using RepeatMasker version 4.1.5^42^. Then, the PacBio Secondary Analysis Tools on Bioconda^36,50^ were used to process the Iso-Seq reads and identify transcripts. Iso-Seq3, a scalable *de novo* isoform discovery from single-molecule PacBio reads workflow was applied on the reads from all five tissue types (liver, gonad, eyes, gills and epipodial tentacle). The full workflow is detailed at https://github.com/ylipacbio/IsoSeq3. Briefly, cDNA primers, polyA tail and artificial concatemers were removed, and de novo isoform-level clustering was performed. High-quality isoforms were mapped to the genome (**Figure 4**) using pbmm2 with a 99.86% mapping rate (samtools-flagstat version 1.16.1^32^). Redundant transcripts were collapsed, and the TAMA^51^ package was used to produce gene models and to identify open reading frames (ORF) and coding regions (CDS). AGAT version 1.2.0^52^ was used to filter all isoforms and to obtain the longest isoform per gene. For functional annotation, the protein-coding genes’ amino acid sequences were blasted (cut-off value 1e^-5^) using (1) blastp^53,54^ against UniProtKB/Swiss-Prot database^55^, (2) KEGG^56,57^, (3) InterProScan version Version 5.59-91.0^58,59^ and (4) eggNOG version 2.1.8^60^ to find protein hits, gene ontology and pathway information. Overall, 25,422 protein-coding genes and 61,149 transcripts were identified. The distribution and content of the gene elements are presented in **Figure 4**. Gene density and methylation density across the 16 pseudo-chromosomes are presented in **Figure 3**.

**Figure 4.**
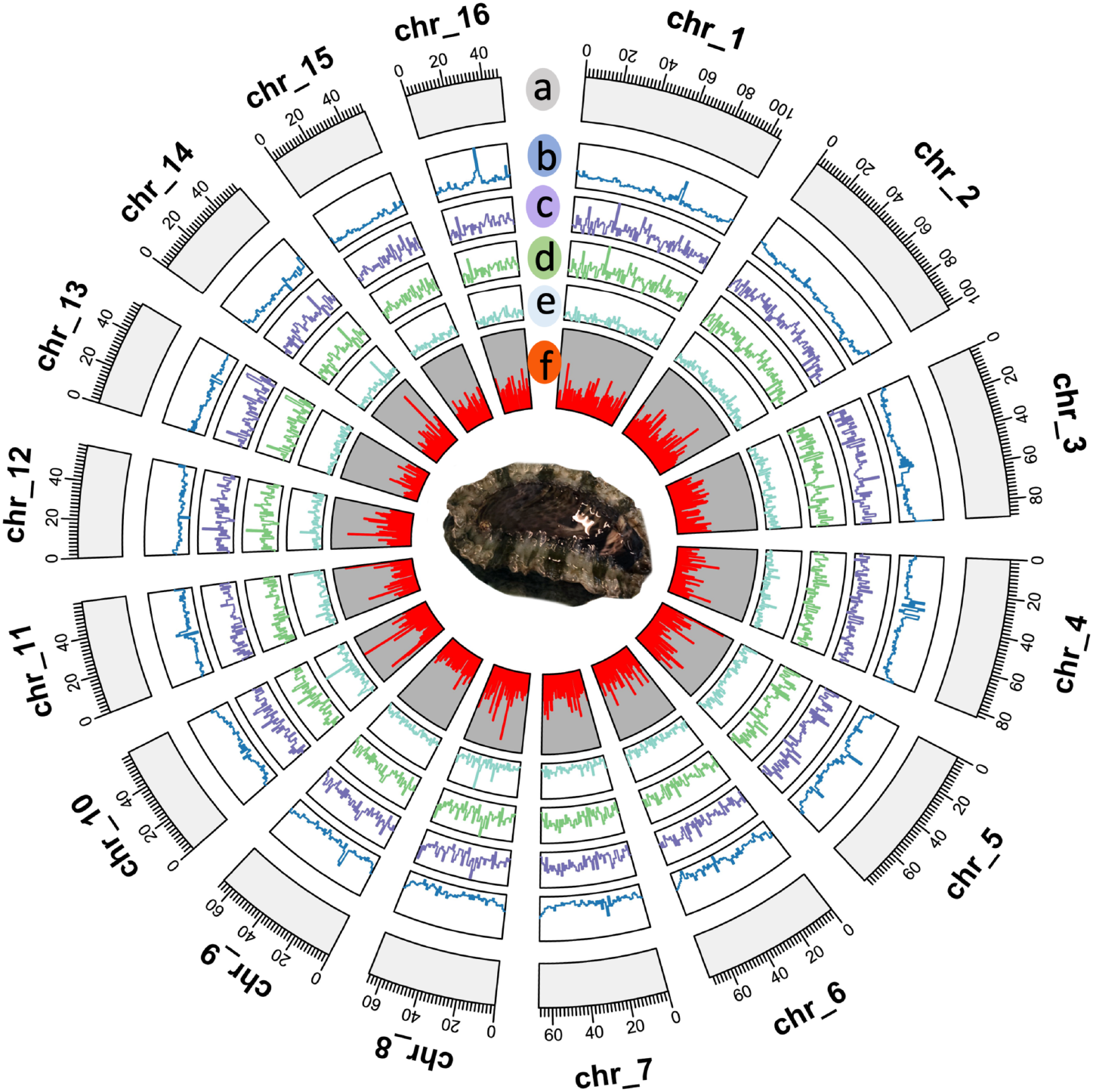
Circos plot of *H. asinina* genome characteristics. From the outer to the inner layer: (a) 16 chromosome-level scaffolds (values represent length in Mb), (b) GC content, (c) exon content, (d) 5’ untranslated region (UTR) content, (e) 3’ UTR content, (f) Iso-Seq mapping coverage and a photo of *H. asinina* female specimen that was used for this study. Generated and calculated using TBtools^80^ on the basis of 1 Mb windows.

### Data Records

The genome assembly and all of the sequencing data used in this study have been submitted to the National Center for Biotechnology Information (NCBI) via BioProject ID PRJNA1080039^61^. PacBio DNA sequencing data is available under the NCBI Sequence Read Archive accession number SRR28083764^62^. PacBio Iso-Seq data for all tissues (liver, gonad, tentacles, gills and eyes) is available under the NCBI Sequence Read Archive accession numbers SRR28084366-SRR28084370^63–67^. The Omni-C data is available under the NCBI Sequence Read Archive accession number SRR28100643^68^. The genome assembly has been deposited at GenBank under the accession GCA_037392515.1^69^. Genome annotation files^70^, repeat sequences files^71^ and the mitochondrial genome assembly^72^ are available in Figshare.

### Technical Validation

#### Nucleic acid

DNA quality and quantity was measured using Thermo Scientific™ NanoDrop (260/280 = 1.87; 260/230 = 2.13, 111.8 ng/μl) and Qubit dsDNA High Sensitivity Assay (106 ng/μl). The integrity of the HMW gDNA was also confirmed by the Australian Genome Research Facility (AGRF) using the Agilent™ FemtoPulse system. RNA quality and quantity from all tissues were measured using Thermo Scientific™ NanoDrop (260/280 = 2.07-2.14; 260/230 = 1.93-2.28) and the Agilent™ TapeStation 4150 system (RIN > 9.3).

#### Sequencing data, assembly and annotations

Using HiFiAdapterFilt version 2.0.0^73^, the PacBio HiFi reads BAM file was converted into a FASTA file prior to the adapter filtering and read trimming (using the default settings). The adapter-free FASTA file was used for k-mer counting using Meryl version 1.4^74^ with *k*=20 (estimated with Meryl based on the genome size). Next, the *k*-mer database was used as input to estimate the overall characteristics of the genome (genome heterozygosity, repeat content, and size) from sequencing reads using a *k*mer-based statistical approach via GenomeScope 2.0 version 1.0.0^75,76^ (**Figure 5**). The Hifiasm primary assembly output was used as input for QUAST version 5.2.0^77^ and Merqury version 1.3^74^ to generate a quality assessment report of the assembly. We used BUSCO version 5.5.0^78^ with the metazoan_odb10 database to assess the genome assembly and annotation completeness, resulting in 97.6% and 93.1% complete BUSCOs, respectively (**Figure *6***). For BUSCO’s annotation completeness (-m prot), isoforms were filtered from the gene set according to the latest BUSCO protocol^79^. Finally, we used Merqury^74^, a reference-free quality and completeness assessment tool for genome assemblies, resulting in 99.54% *k-mer* completeness and an assembly consensus quality value (QV) of 65.5 (>99.99% accuracy).

**Figure 5.**
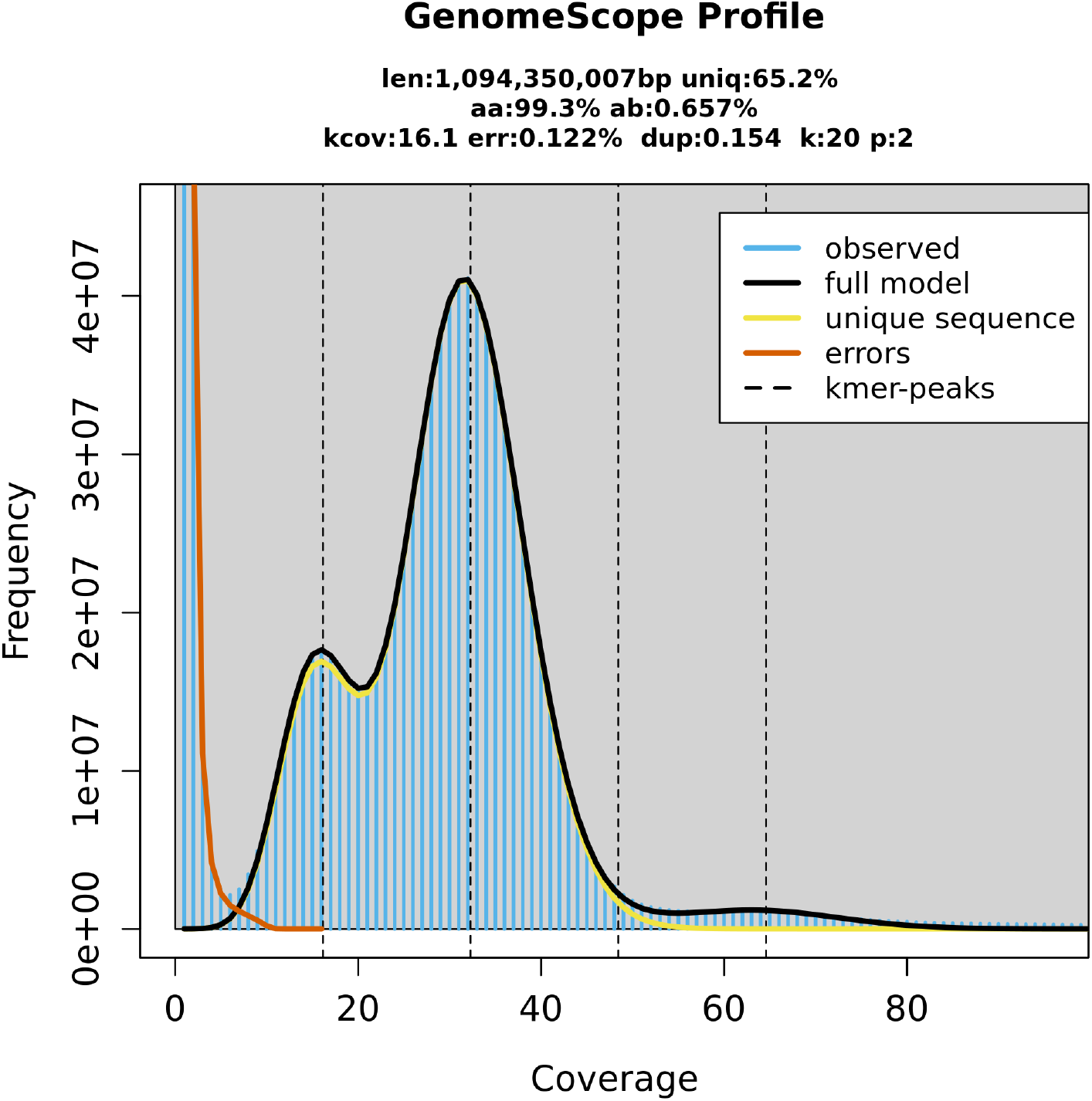
GenomeScope profile of *H. asinina* genome. Len: estimated genome length; uniq: overall length of unique, non-repetitive sequences; het: heterozygosity rate; kcov: mean k-mer coverage for first peak; err: error rate; dup: read duplication rate; k: k-mer length (automatically assigned).

**Figure 6.**
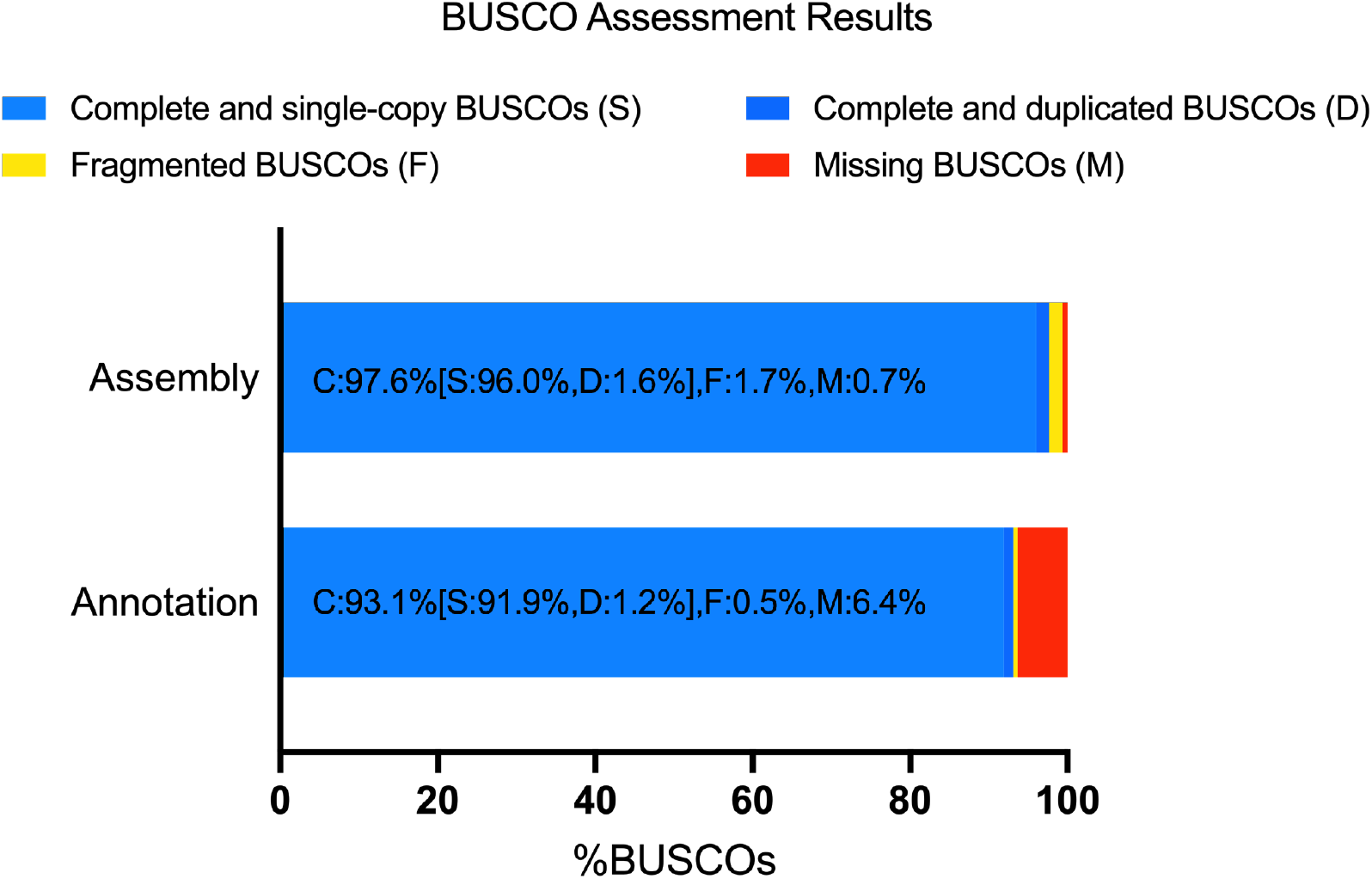
BUSCO assessment results. BUSCO assembly (--genome) and annotation (--prot) assessment based on metazoa_odb10 lineage dataset (number of genomes: 65, number of BUSCOs: 954).

## Code Availability

Except where otherwise stated, bioinformatics tools and software were used with default parameters, and all code used for this assembly can be found at https://github.com/roybarkan2020/AbsGenome. In addition, a list of the tools and software used for the assembly is provided in the Methods section (with references to the tool publication, which includes a link to the tool manual and/or GitHub link).

## Acknowledgements

The authors would like to acknowledge the Marine and Aquaculture Research Facility (MARF, James Cook University, Townsville) team, Cairns Marine (Cairns, Queensland, Australia) for collecting the abalone, the Australian Genome Research Facility (AGRF) team – Dr Dhanya Sooraj, Trent Peters and Saurabh Shrivastava. We would also like to acknowledge Dr Inga A. Frøland Steindal, Dr Bruna Louise Pereira Luz and Julia Yun-Hsuan Hung for their lab support.

## Author contributions

Study design: R.B., J.S., I.C. and S.A.W. Laboratory work: R.B. and S.C.Y.L. Data analysis and interpretation: R.B., J.S., I.C., and S.C.Y.L. Drafting the manuscript: R.B., J.S., I.C., S.A.W and S.C.Y.L.

## Competing interests

The authors declare no competing interests.

## Notes

https://www.ncbi.nlm.nih.gov/bioproject/PRJNA1080039

https://www.ncbi.nlm.nih.gov/datasets/genome/GCA_037392515.1/

https://figshare.com/s/696aef84d9e128b201e6

https://figshare.com/s/b8d6aa4462d2bdf048ee

https://figshare.com/s/6c6f99dd19ed64286105

### Competing Interest Statement

The authors have declared no competing interest.

